# The Cost of Appearing Suspicious? Information Gathering in Trust Decisions

**DOI:** 10.1101/495697

**Authors:** I. Ma, A. G. Sanfey, W.J. Ma

## Abstract

Trust decisions are inherently uncertain, as people have incomplete information about the trustworthiness of the other prior to their decision. Therefore, it is beneficial to gather information about a trustee’s past behaviour before deciding whether or not to trust them. However, elaborate inquiries about a trustee’s behavior may change the trustee’s willingness to reciprocate, causing either a decrease due to the investor appearing suspicious, or an increase because the investor appears to be highly betrayal-averse. Such a change could cause the investor to gather less or more information, respectively. We examine how information acquisition is modulated by social context, monetary cost, and the trustee’s trustworthiness. Participants had the opportunity to sequentially sample information about a trustee’s reciprocation history before they decided whether or not to invest. On some trials, we induced a social context by telling the participant that the trustee would later learn how much the participant sampled (“overt sampling”). Participants sampled less when there was a monetary cost and when the reciprocation history was more conclusive. Crucially, when sampling was free and overt, participants sampled less, suggesting negative consequences of appearing suspicious. In post-experiment questionnaires, participants indeed reported a belief that the reciprocation probability would decrease when information was overtly sampled. The findings replicated in a second experiment and were well accounted for by a utility-maximizing model in which overt sampling induces a decrease in reciprocation probability. This study opens the door to broader applications of the tools and models of information sampling to social decision-making.

**Significance Statement:** Trust and reciprocity are essential for establishing and maintaining beneficial cooperative interactions. However, not everyone can be trusted. Here, we focus on a fundamental question in the study of social interaction: how people gather and use information to make a decision to trust or not trust. While more information seems better, participants gathered less information about trustworthiness when the trustee would learn about the inquiries, as if they avoided appearing suspicious. Indeed, participants reported that they believed that sampling elaborately would make the trustee less willing to reciprocate trust. Using a mathematical model of information gathering, we show that this belief indeed reduces the value of information. Our findings contribute to a deeper understanding of information gathering in social contexts.

---

The processes of trust and reciprocity are essential for establishing and maintaining beneficial human cooperative interactions^1^. When trust is invested, and then reciprocated in turn, the investor (who makes the decision to trust) and the trustee (in whom trust is placed) typically emerge better off from the exchange. However, a trustee can often maximize their own profit by not reciprocating the trust placed in them. Therefore, perhaps the key factor in a successful cooperative interchange is the investor’s belief about whether or not the trustee will reciprocate. Understanding how an investor gathers and uses information to make a decision to trust or not trust is a fundamental question in the study of social interaction.

The possibility that one’s trust might be violated confers inherent uncertainty to the decision of whether or not to trust. Some individuals are more trustworthy than others, and repeatedly interacting with the same person does not necessarily result in the same outcome^1,2^. In addition, we often only have limited information about someone’s level of trustworthiness. Therefore, decisions that involve trust are both risky and uncertain: there is variability in the outcome given a specific degree of trustworthiness of the other person (risk), and there is incomplete knowledge about this trustworthiness (uncertainty, sometimes also referred to as ambiguity). Due to this uncertainty, it is beneficial to acquire information to determine whether someone can be trusted, which can be done by gathering information about the trustee’s past behaviour^3^. However, there are specific consequences to information sampling in a social context; your inquiries might affect the other person’s impression of you^4,5^. On the one hand, if someone continually checks up on our reliability, it may make us less likely to behave pro-socially with that person in the future, for example because too many detailed questions could be seen as offensive. As a consequence, the investor should gather less information when the trustee is aware of the inquiries than when they are not. On the other hand, extensive information search is typically indicative of deliberation or caution, which may be warranted if the decision is risky^6^. Therefore, the investor could alternatively attempt to communicate a strong aversion to betrayal by sampling *more* when the trustee is informed. This may be beneficial because reducing uncertainty about how a self-serving choice negatively impacts the well-being of another is shown to increase prosocial behavior, especially in those with high empathy^7^. Therefore, if the trustee realizes that the investor is highly betrayal-averse as indicated by extensive sampling prior to trusting, then the trustee may become more likely to reciprocate, especially if the trustee is highly empathic. Given this mechanism, the investor should gather more information when the trustee is aware of the inquiries. In view of these opposing predictions – sampling more to communicate betrayal aversion or sampling less to avoid appearing suspicious – the goal of our study is to understand how people use information sampling to signal their motives in building trust.

To this end, we designed a novel version of a single-shot trust game, the Information Sampling Trust game (IST). This game allowed us to investigate information sampling in trust decisions and, within subjects, compare between overt (trustee is informed of sampling) and covert (trustee is not informed of sampling) information acquisition. Each participant completed the IST in the role of investor and was given the opportunity to sequentially gather information about a trustee’s previous reciprocation history before deciding whether or not to invest (to trust) (Figure 1). We told the participant that the trustee would be different on each trial. Unknown to the participant, each trustee was computer-generated; we drew the trustee’s reciprocation probability *r* pseudo-randomly from six possible values (0, 0.2, 0.4, 0.6, 0.8, and 1). The trustee’s revealed past choices were independent draws from a Bernoulli distribution with parameter r. When the participant decided to finish sampling, they chose whether or not to invest. The experiment consisted of four conditions: a monetary cost of sampling (either costly or cost-free), crossed with social context (overt or covert). We told the participant in the instruction phase that after the experiment, we would randomly select three trials. For the subset of the three trials on which the participant had decided to not invest, the participant’s earnings would be equal to their endowment. For the subset of the three trials on which sampling was overt and the participant had decided to invest, we would contact the corresponding trustees and tell them how many samples the participant had drawn; next, the trustees would decide to either reciprocate trust by returning 50% of the multiplied endowment to the investor, or defect/betray, which meant keeping all the money, so that the participant would end up with €0. In the *Covert* condition, the participant was told that the trustee would not be informed of the information sampling and instead, the trustee believed that the decision to invest was not based on any prior information. Participants were not instructed about any potential effects that the overt and covert conditions might have. We told the participant that their final pay-out would be the average of their earnings on the three selected trials. In reality, we followed this procedure except that the trustees’ decisions were simulated using their respective reciprocation probabilities.

**Figure 1.**
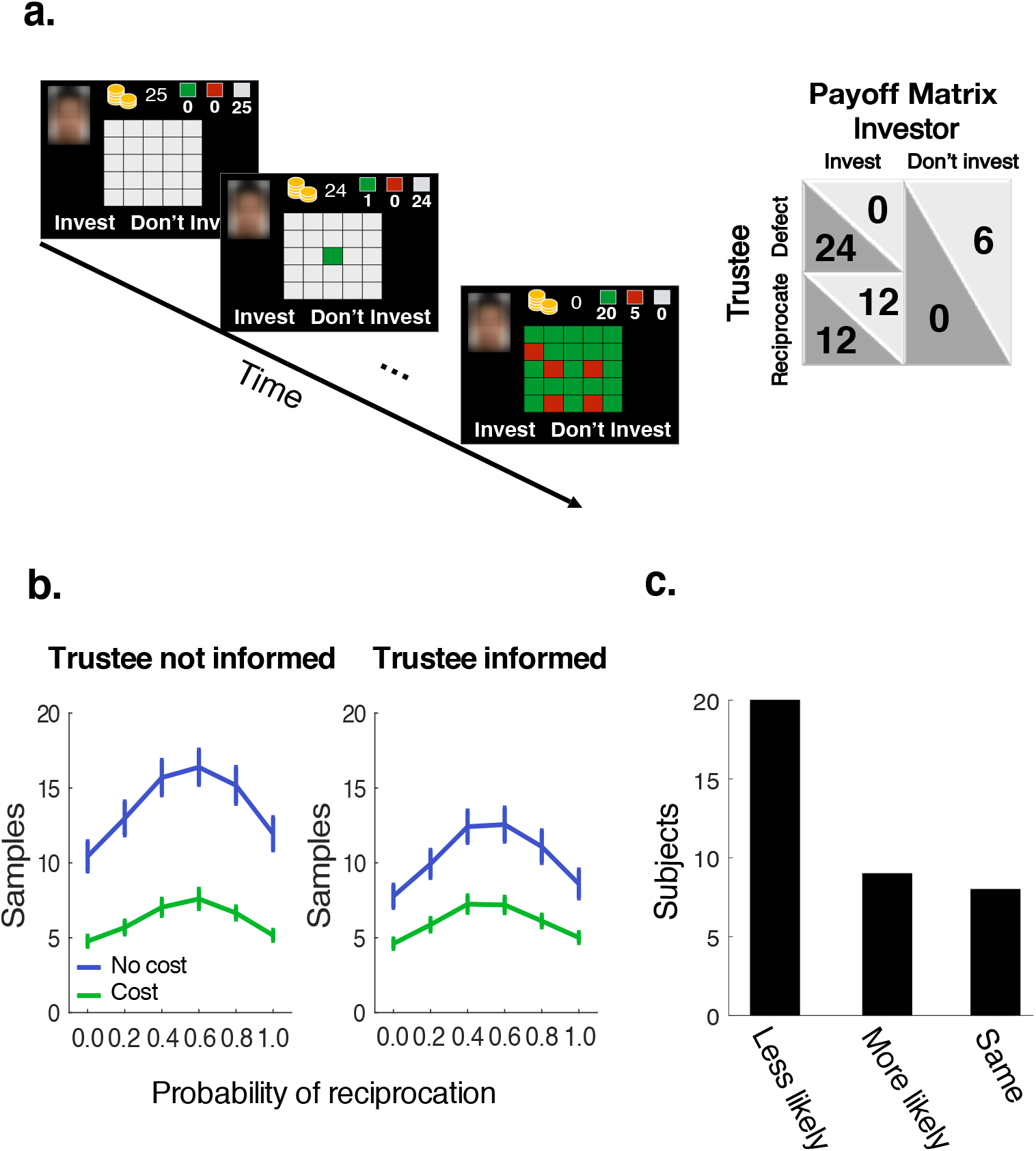
Task and data. **A.** Trial sequence and payoff matrix in the Information Sampling Trust game (IST). Before the participant makes an investment decision, they could sequentially sample a trustee’s reciprocation history with other investors. On each trial, information could be sampled up to 25 times. The colour of the opened box indicated the trustee’s past decision. Green = reciprocated trust, red = did not reciprocate trust, grey = not sampled. In the monetary cost conditions, 5 eurocents were deducted for each decision to sample (from a separate sampling budget of 125 eurocents per trial). **B.** Data which shows the effect of monetary cost and social context. Mean and SEM of drawn samples for each generative reciprocation probability per cost condition: green = monetary cost, blue = free. Left panel = trustee not informed, right panel = trustee informed. **C.** Beliefs about the consequence of overt sampling. The most common belief amongst participants was that the more information they overtly sampled, the less likely reciprocation would become. Participants could choose from one of 3 categories: overtly sampling more information would make reciprocation less likely, more likely, or stay the same.

## RESULTS

We assessed the effects of monetary cost, social context, and outcome uncertainty (the distance between the reciprocation probability and 0.5, where *r* = 0.5 is the maximum outcome uncertainty) on the decision to sample or stop, using a mixed-effects model. There was a significant interaction between monetary cost and social context (coefficients mean±SEM; *β* = −0.10±0.01, *p* < 0.001); participants sampled less if sampling was overt and monetarily cost-free (*β*= −0.45±0.04, *p* < 0.001) but not when samples were monetarily costly (*β*= 0.04±0.04, *p* = 0.37). Participants also sampled less when sampling was monetarily costly compared to cost-free, in both overt (*β* = −0.64±0.04, *p* < 0.001) and covert conditions (*β*= −1.06±0.04, *p* <.001). As expected, outcome uncertainty also affected sampling (*β*= 0.41±0.01, *p* < 0.001). Specifically, information sampling increased when outcome uncertainty was higher, and the acquired information was thus relatively inconclusive (as would be the case when *r* is close to 0.5). 1b). Specifically, we further examined this with Bonferroni-corrected pairwise t-tests, which revealed that people sampled more when *r* = 0.8 than when *r* = 0.2 (*t*(40) = −2.05, *p* < 0.001, *d* = 0.59) but not for *r* = 0.4 compared to *r* = 0.6 (*t*(40) = −1.44, *p* = 0.010, *d* = 0.42) or for *r* = 0 compared with *r* = 1 (*t*(40) = −0.88, *p* = 0.085, *d* = 0.28).

We then used a separate logistic regression to test whether the decision to invest was predicted by *r* and the conditions. This logistic regression returned a coefficient *β* = 9.12±0.18 (*p* < 0.001) for *r*, indicating that the probability of investing increased with a higher *r*. This confirms that the acquired information was used in the investment decisions. As expected, monetary cost, social context and their interaction were not significant predictors of the decision to invest (monetary cost: *β*= −0.023±0.095, *p* = 0.81; social context: *β*= 0.011±0.096, *p* = 0.91; interaction between monetary cost and social context: *β* = 0.113±0.135, *p* = 0.40). After task completion, participants were asked to indicate whether they believed that when sampling was overt to the trustee, more sampling made reciprocation more likely, less likely, or did not change the probability of reciprocation. Believing that overt information sampling would make reciprocation less likely was the most commonly reported answer (test for non-uniformity: *χ*^2^(2, *n* = 37) = 19.28, *p* < 0.001, Figure 1c). This suggests a cost of appearing suspicious and is consistent with the intuition that if someone continually checks up on our reliability, it may make us less likely to behave pro-socially with that person in the future.

### Computational models of information sampling

Based on the behavioural data alone, it is not possible to know if this belief about the consequences of appearing suspicious can quantitatively account for the difference between the overt and covert sampling conditions. We therefore developed an normative sampling model under the assumption that every sample reduces the reciprocation probability by a constant factor. We refer to this normative model as the *Cost of Appearing Suspicious* (CAS) model (see Methods). The model is Bayesian in the sense that the agent computes a posterior belief distribution over another’s trustworthiness. The agent uses this posterior to calculate for every possible state in the task which action has the highest expected utility: sampling or stopping (to invest, or not invest). We derived these expected values of all state-action pairs using the Bellman equations and dynamic programming^8^. In the overt sampling conditions, the value of investing takes into account the factor *w* by which the agent believes the trustee’s reciprocation probability will decrease with each sample drawn. In the monetary cost conditions, the model accounts for the immediate subjective cost of sampling, c. We allow for two deviations from optimality as suggested by the (repeated) trust game literature: subjective prior beliefs^9^ and risk aversion (often referred to as betrayal aversion in the context of trust)^10^. The CAS model improved in fit to the data when these parameters were added (Table S1). In simulating this forward-looking model, we find that lowering *ω* has the consequence of decreasing the utility of sampling at any point, which, in turn, decreases the agent’s likelihood of sampling.

We find that the CAS model fits well (see Figure 2a for the data per state and the predicted data by the CAS model, see Figure 2b for the model fit on the summary statistics). In particular, the CAS model accounts for the difference between the overt and covert sampling conditions. This result suggests that the participants’ self-reported beliefs about the consequences of appearing suspicious underlie the effect of social context on their behavior.

**Figure 2.**
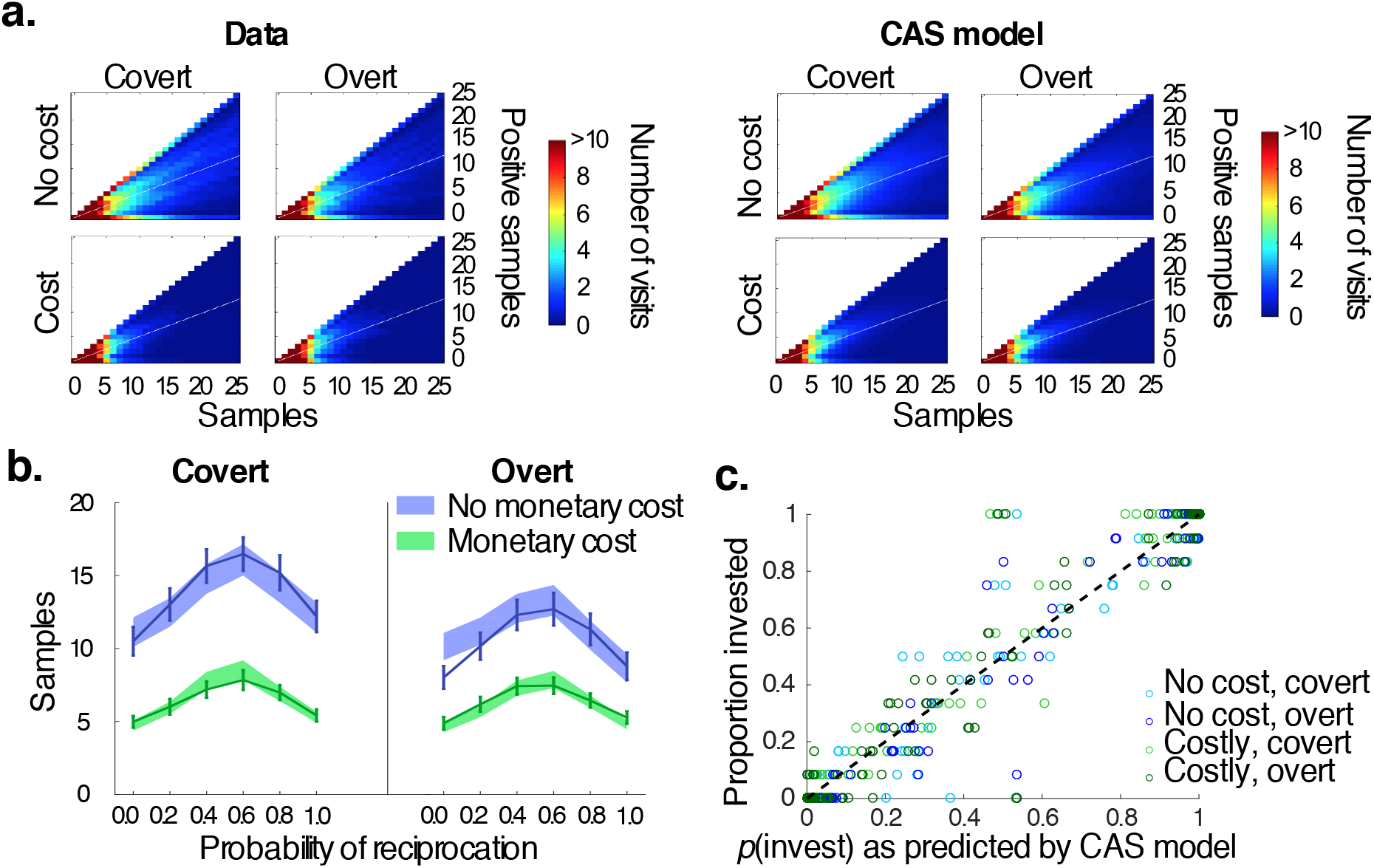
Model fits to the sampling decisions for each condition. **A**. Left pandel is the number of times that each possible state was visited in the behavioral data. Right panel shows the number of times that each possible state was visited as predicted by the fitted CAS model (synthetic data). Synthetic data was generated by the CAS model using estimated parameters, averaged over participants for each of the four conditions. The full state space is determined by every possible combination between the number of samples (x-axis) and number of positive samples (y-axis). Note that reciprocation probabilities of 0 and 1 follow one specific path in the state space, whereas other values of *r* “spread out” in the middle due to stochasticity in the outcome of a sample. Therefore, trials in which participants reached the states along the upper edge (*r* = 1) and bottom edge (*r* = 0) paths occur in high frequency, even though participants sampled more when *r* was closer to 0.5 (as depicted in plot a; white line through the middle is included as a reference). These plots highlight that CAS fits well. **B**. Number of samples per condition and CAS model fit. The average number of samples across all subjects is represented by lines (mean) and error bars (SEM). Shaded areas indicate SEM of the model fit. **C**. The CAS model fits to the investment decisions per condition for each participant. The diagonal (in black) represents a perfect fit. Monetary cost conditions in green, monetarily free in blue. Lighter colors reflects the conditions in which sampling was covert. The CAS model predicts the investment decisions well.

We tested if participants’ behavior was better fitted by heuristic models than the normative model (for formal descriptions of the heuristic models see SI). The heuristic models are less motivated by general principles and more specific to the task at hand. Therefore, we do not a priori favour these models. The *Sample cost* model is similar to the CAS model, except that the agent does not believe sampling will decrease probability of reciprocation. Instead, in this model the cost of a sample is dependent on the condition (i.e., four free cost parameters, see SI). The implementation of forward-reasoning models like the CAS model and the Sample cost model is likely computationally expensive and the brain might instead use a “good enough”, simpler strategy^11,12^. We therefore also examined computationally simpler strategies. In the *Uncertainty model,* the agent continues sampling until uncertainty about trustworthiness – as measured by the width of the posterior belief distribution – drops below a level that they find tolerable; this tolerance may vary between conditions (SI). The *discrete Drift Diffusion Model* (dDDM) has an intuition similar to that of the standard Drift Diffusion Model^13^. Here, we consider the hypothesis that people do not use Bayesian posterior beliefs but instead maintain criteria for when they view a trustee’s behaviour as trustworthy or untrustworthy, and sample until the evidence meets one of those criteria. This requires keeping track of the sample outcomes in favour of investing and not investing. The decision to stop sampling information is then determined by whether their difference is sufficiently large, i.e., when the difference reaches a bound b. We allowed the bound to vary between the four conditions (SI). It should be noted that the dDDM is not equivalent to the DDMs that are often used in perception studies^14^. First, in perception studies, the noise is typically Gaussian internal noise, whereas here, it is Bernoulli noise associated with past investment outcomes. Second, in perceptual applications, the time scale is hundreds of milliseconds to seconds, whereas here, accumulation takes place over a much longer time scale (tens of seconds). Finally, accumulation of evidence in regular DDMs is passive, whereas the dDDM describes a process in which the agent makes a decision at every time step.

We tested different versions of each model (SI). For the CAS model and Sample cost model, these included a risk attitude term and a prior belief. For the Uncertainty model, we included a prior belief, and for the dDDM, we tested versions with assymetric bounds and collapsing bounds (see SI). We then selected the winning version of each model and compared those with each other. Models were fitted to the data at the individual level using a log likelihood optimization algorithm as implemented in the fmincon routine in MATLAB (©Mathwork). The optimization was iterated 100 times with varying initiations to avoid local minima.

Behavior was not fitted better by the heuristic models than by the CAS model (Table 1). Of the heuristic models, the Uncertainty model fitted best and approximately as well as the CAS model. Here, we see a tension between interpretability and goodness of fit; the heuristic models are more concerned with fitting the data and less with psychological content^15^. Specifically, the Uncertainty model leaves open the question through what process people change their uncertainty criterion in social contexts. By contrast, the CAS model is based on a psychologically motivated mechanism and might therefore be more generalizable to other studies that examine how people change their information search when they are being monitored^4,16,17^. In light of these considerations, we focus on the CAS model from now on.

**Table 1.** Model comparisons between CAS and each heuristic model Study 1

|  | **AIC** |  |  | **BIC** |  |  |
| --- | --- | --- | --- | --- | --- | --- |
|  |  | 95% CI |  |  | 95% CI |  |
|  | median | LB | UB | median | LB | UB |
| CAS vs. Sample Cost | -825 | -70 | -1610 | -825 | -109 | -1725 |
| CAS vs. Uncertainty | 1268 | -219 | 2326 | 1681 | 444 | 2949 |
| CAS vs. dDDM | -4786 | -6175 | -3849 | -3960 | -5224 | -2787 |

95% CI = Bootstrapped 95% confidence interval of the summed difference between model fits. LB = lower bound, UB = upper bound. Each heuristic model's evidence was subtracted from that of CAS. Smaller AIC and BIC values indicate better fit. Thus, negative values indicate a better fit for CAS. Because of the summation of the difference, large positive or negative numbers therefore indicate that one model wins for most subjects.

CAS parameter estimates confirmed that the suspicion-induced decrease in reciprocation probability *ω* was stronger when there was no monetary cost of sampling (median = 0.9998, 95% CI [0.9985, 0.9999) then when there was a monetary cost of sampling (median = 0.9999 95% CI [0.9993, 1.000], Wilcoxon signed-rank test: *Z* = −2.07, *p* = 0.038). The cost-of-sampling parameter was estimated higher in the monetary cost condition (median = 0.512, 95% CI [0.049, 1.013]) than in the no-cost condition (median = 0.00003, 95% CI [0.000, 0.0008], Wilcoxon signed-rank test: *Z* = −5.04, *p* < 0.001). On average, people showed mild risk aversion (median risk attitude = 0.116, 95% CI [0.039, 0.167]; see Table S2 for parameter estimates of the heuristic models).

### The model predictions of invest decisions

Next, we examined whether the CAS model could predict investment decisions (after sampling has concluded) by using the parameter estimates obtained from fitting the choices from the sampling stage. To do so, we extracted the expected utility of investing at the time of stopping from the CAS model. We used this utility to predict investment decisions, using a softmax model with a bias and a decision noise temperature (equation 9). The CAS model fitted well to the investment decisions (*β* = 8.573, *p* < 0.001, Nagelkerke pseudo-*R*^2^ = 0.643; Figure 2c). No other model performed substantially better: the 95% CI of the summed difference in log likelihoods for the alternative models relative to the CAS model was [-59, 533] for the Uncertainty model, [-57, 540] for the Sample cost model, and [-651, 289] for the dDDM.

### Study 2: replication in biased generative reciprocation probabilites

We conducted a second study to test the robustness of the “cost of appearing suspicious” effect under variations of the distribution of the reciprocation probability (n = 75; see SI). The experimental procedure was identical to Study 1, apart from the fact that participants sampled over either positively biased (r = 0.2, 0.4, 0.6, 0.8, and 1), or negatively biased (r = 0, 0.2, 0.4, 0.6, and 0.8) reciprocation probabilities. Our findings of social context and monetary cost replicated and the heuristic models again did not fit better than the CAS model (Table S5). Using a mixed model we showed an interaction effect between social context and monetary cost (*β* = −0.059 ±0.014, *p* < 0.0 01): people sampled less when it was overt to the trustee and sampling was cheap (*β* = −0.248 ±0.041, *p* < 0.001) but not when sampling was monetarily costly (*β* = −0.013±0.039, *p* = 0.734). People sampled more when sampling was monetarily free compared to costly both during covert sampling (*β* = −0.866±0.040, *p* < 0.001), and overt sampling (*β* = −0.630±0.040,*p* < 0.001). They also sampled more when outcomes were inconclusive (i.e., high outcome uncertainty main effect *β* = 0.875±0.014, *p* < 0.001) but there was an interaction between bias and outcome uncertainty (*β* = 0.1279±0.014, *p* < 0.001) as the effect of outcome uncertainty was significant in both groups yet stronger in the positively biased (*β* = 1.069±0.044, *p* < 0.001) compared to the negatively biased groups (*β* = 0.753±0.033, *p* < 0.001). There were no or other significant interactions (all *p* > 0.256). The comparisons between different versions of the models replicated for the all models except for the dDDM; asymmetric bounds improved the model fit in Study 2. Heuristic models again did not fit better than the CAS model (table S5).

## DISCUSSION AND CONCLUSIONS

Acquiring information to reduce our uncertainty about the intentions and future actions of others is crucial to determine our social behavior. Understanding why and about whom we gather information has been the objective of many studies in social psychology (for review see^18^). However, one often ignored aspect of social information sampling is that people might take into account the fact that the other is aware of the inquiries and adjust their sampling accordingly. Here, we examine how trustworthiness information sampling changes when it is overt. Across two studies, we demonstrated a cost of appearing suspicious effect: people discounted the value of investing when the quantity of acquired information was overt to the trustee. Although we did not instruct our participants about whether, or how, information sampling might affect reciprocation probabilities, most subjects reported that they believed that sampling would decrease the reciprocation probability. We developed the normative CAS model based on this stated belief. In this model, the belief that sampling will reduce the reciprocation probability directly causes a decrease in the utility of sampling, which is consistent with the behavior in the task.

The central concept of the CAS model is closely related to behavior in other situations when they are being monitored^16^. Previous studies have shown that people care about the intentions of others in cooperative decisions^19,20^. Information gathering can reveal the investors’ covert intentions that govern the cooperative decision-making process. For example, it has been proposed that some individuals care about whether their cooperation partner gathers information about their own profit if they were to defect, before making a cooperative decision that is mutually beneficial^16,17,21^. Under that assumption, simulations of cooperative games show that blind cooperation becomes a beneficial strategy^16,17,21^. For example, when individuals compete to become cooperative partners in heterogeneous populations with both reliable and unreliable individuals, those individuals who blindly cooperate should be preferred^21^. The assumption of these simulations is corroborated by behavioural experiments showing that people who gather information about their own profit are perceived as more selfish, and indeed also tend to behave more selfishly^5^. This converging evidence suggests that cooperative decision-making incorporates a cost of appearing selfish when assessing the potential payoff structure. Here, we extended this concept to the cost of appearing suspicious by sampling interpersonal information, that is, sampling information about the trustworthiness of others.

Independent of the cost of appearing suspicious, participants were clearly sensitive to outcome uncertainty. People gathered most information when *r* was close to 0.5, and least information when *r* was deterministic. While this is qualitatively optimal, participants also showed an interesting valence-dependent asymmetry in information gathering. Specifically, when outcomes were stochastic, people sampled more when outcomes were mostly positive and less when outcomes were mostly negative. This asymmetry in sampling indicates that beliefs about untrustworthiness are updated faster than beliefs about trustworthiness. This is a similar mechanism as that reported in sampling from experience, where it is more likely that early negative samples cause the agent to stop sampling and avoid someone, even if those samples were not representative^22^. This asymmetry in sampling is also broadly consistent with well-established findings in impression formation and person perception, in which negative behaviour is generally viewed as more diagnostic of morality than positive behaviour^23–26^. However, the mechanism proposed for those findings – that good people more often make selfish decisions than bad people make generous decisions – is not the mechanisms we propose here; instead, we explain the asymmetry through risk aversion: investing is a realistic option but a risky one that requires more information.

A potential negative consequence of sampling less to avoid appearing suspicious is that the subsequent investment decisions are less well informed. In particular, prior beliefs about someone’s trustworthiness based on facial judgement of trustworthiness, attractiveness, narratives about moral character, and previous social experiences in unrelated settings, can bias learning about trustworthiness^9,27–29^. If an initial sample is consistent with the prior information such as trustworthiness judgements based on attractiveness, then it is more likely that the investor will stop sampling even if the information is not representative of the actual trustworthiness. Given that avoiding to appear suspicious reduces information search, the subsequent decision to trust or not might become even more susceptible to these prior beliefs. Future studies are required to test if avoidance of appearing suspicious will indeed make people more sensitive to prior information, especially prior information that has very little or highly uncertain diagnostic value, such as facial attractiveness.

We told subjects that the trustee would be informed of the sampling. In many daily-life scenarios (e.g. gossip), people are uncertain that the trustee will learn about the inquiries. Future studies could inform the trustee about sampling behaviour in a stochastic manner. The model is straightforwardly extended to this situation: the utility of investing will become a linear combination of two terms. Another potential extension could involve directly probing trustee behaviour. Understanding whether and how trustees change their behaviour when investors gather information is important in understanding whether avoiding to appear suspicious is truly adaptive.

Our findings might provide a benchmark to uncover social value computation aberrations in psychiatric disorders. Some psychiatric disorders can be characterized by biases in information sampling, such as insufficient information gathering in addiction^30^, asymmetric weighting of negative evidence in depression^31^, and impaired information sampling cost signals associated with compulsivity^32,33^. Importantly, specific impairments in the ability to model the moral character of others or to respond to social signals are central to a range of psychiatric disorders, including borderline personality disorder and autism spectrum disorder^34,35^. For example, modelling people’s modelling of another person’s state of mind can identify maladaptive social behaviours^36^. More generally speaking, our study opens the door to broader applications of the tools and models from information sampling to understand social decision-making.

## METHODS

### Study 1

#### Participants and experiment procedure

A total of 41 participants without colorblindness (13 males) completed the experiment (age *m* = 22.95, *sd* = 3.71, range = 18-34). Three participants misunderstood instructions and were excluded. Data from one participant was excluded due to a technical malfunction. The data were collected in the behavioural labs of the Donders Institute, Nijmegen, the Netherlands. The study conformed to the Declaration of Helsinki and was approved by the local ethics committee (CMO2014/288). Written informed consent was obtained from all participants before study enrolment. In the *no cost* condition, there was no monetary cost of sampling. Each participant completed 60 trials in each of the four conditions, with the four sets of trials being intermixed in random order. The task therefore consisted of 240 trials in total, resulting in a maximum of 6000 sampling decisions, and took approximately 40 minutes to complete. On *no cost* trials, the entire sampling budget (125 euro cents) was added to the trial outcome, while on *cost* trials, the number of opened boxes times 5 euro cents was subtracted from the sampling budget and the remainder then added to the trial outcome.

### Computational Models

#### General structure of models

The state is determined by the number of open green boxes, *n*_+_, and the number of open red boxes *n*_-_. We consider four models, all with their own variants: A Cost of Appearing Suspicious (CAS) model, a Sample cost model, an Uncertainty model, and a discrete Drift Diffusion model (dDDM) (see SI for formal description of the heuristic models). In each model, the agent computes a decision variable based on *n*_+_ and *n*_-_, and the probability to stop sampling is a noisy function of this decision variable. After the agent stops sampling, the same decision variable determines the probability that the agent invests. The CAS model, the Sample cost model, and the Uncertainty model are based on an evolving posterior distribution over the trustee’s reciprocation probability *r*. Through parameter recovery and model recovery, we verified that the number of decisions was large enough to accurately estimate parameters and that the models were distuinguishable (see SI).

#### Cost of Appearing Suspicious (CAS) model

The CAS model is based on the agent computing expected utility through forward reasoning. It consists of four components: prior beliefs over reciprocation probability *r*, an evolving posterior distribution over *r*, iterative maximization of future expected utility under this distribution, and decision noise. We assume that the prior over *r* is a beta distribution with parameters *α*_0_ and *β*_0_. The posterior over *r* is a beta distribution with parameters *α* = *n*_+_ + *α*_0_ and *β* = *n*_-_ + *β*_0_. An investment outcome can have two values: reciprocation (outcome = 1, then the investment amount is multiplied by *m* = 2) or betrayal (outcome = 0, then the investment amount is multiplied by *m* = 0). The investment outcome follows a Bernoulli distribution with parameter *r*, i.e *p*(outcome=1|*r*) = *r*. When building a belief about the outcome of an investment decision, the agent does not know *r* and therefore has to marginalize over *r* using the current posterior over *r*, which isp(*r*|*α, β*). This gives the conditional distribution of outcome given *α* and *β*:

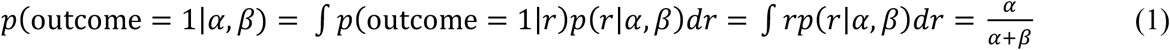

and:

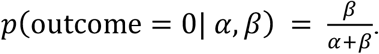

This is a distribution with mean 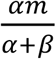 and variance 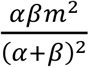.

We hypothesize that the agent believes that overtly sampling induces a decrease in the reciprocation probability *r*. Since people sample over the trustee’s *past* decisions, such a belief does not affect the probability of a positive *sample* outcome. However, if the investor invests after terminating their sampling, the reciprocation probability that the investor believes they will experience is not the *r* from the trustee’s past (the one that the investor has worked to infer through sampling), but a modulated *r*, where every sample that the investor has drawn multiplies *r* by a factor ω that takes values between 0 and 1. Thus, if the investor has drawn *n* samples, the reciprocation probability is:

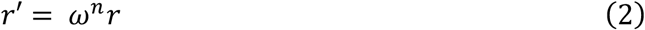

The posterior distribution over *r*’ is now a rescaled and truncated beta distribution. The distribution *p*(outcome|*α,β*) now has a mean of 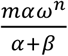 and a variance of 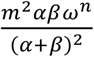. The expected utility of investing gets modified accordingly: We fit one ω for each of the two overt condition (when the trustee is informed of the sampling), that is, once for overt sampling without a monetary cost, and once overt sampling with a monetary cost. The weight parameter *ω* is 1 when sampling is covert sampling (i.e., when the trustee is not aware of the sampling).

We are now ready to define the expected utility of not investing and investing. The expected utility of not trusting is *U*_0_ = 1, the normalized endowment, which is independent of *α* and *β*. The expected utility of trusting should contain the expected amount earned from an investment. In addition, people are also known to differ in their risk attitude. Specifically in trust games, people may be betrayal-averse ^37^ To model such a risk attitude, we subtract a multiplier times the variance of the amount earned from the expected amount earned. The overall expected utility of trusting, which we denote by *U*_1_, is weighted by the decrease in *r*, and thus becomes:

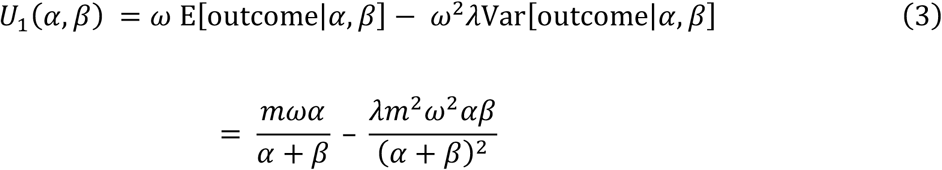

where *λ* parametrizes risk attitude, with *λ*>0 representing risk aversion and λ<0 risk-seeking.

At any time, the participant has two possible actions: stopping (*a* = 0) and sampling (*a* = 1), except when *t* = *T*+1, when only *a* = 0 is available. The expected value of a state-action pair, *Q*(*a, β*; action), is given by the Bellman equations^38^. Specifically, if at time *t*, the participant stops (*a* = 0), then the expected value of the state (*α, β*) is the higher of the expected utilities of not investing and investing:

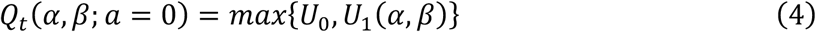

When *t* = *T*+1, the value of the state (*α, β*) is *V*_*T*+1_(*α,β*) = *Q*_*T*+1_(*α,β*; 0). At earlier times, *V_t_* is the larger of the two expected utilities:

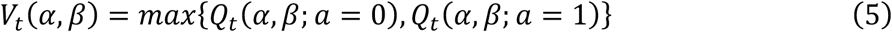

The expected value of a sampling action at time *t* in the state *α, β* by *Q_t_*(*α,β;,a* = 1) is:

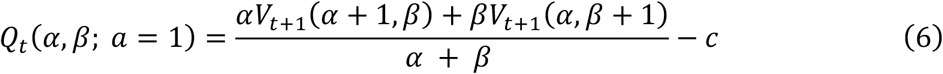

where *c* is the subjective cost of a sample. We fit one cost *c* for each monetary cost condition, i.e., once for when it is monetarily cost free to sample, and once when sampling is monetarily costly (irrespective of overt or covert sampling).

By starting with the final state (when *n* = *T*) we can apply the equation for the expected utility of stopping (equation 4 for *t* = *T*+1, *Q_t_*(*α,β;, a* = 0)). We can then work our way back in time, to obtain the optimal solution for every possible state (dynamic programming^8^). The decision variable, denoted by DV, is the difference between the utilities of sampling and stopping:

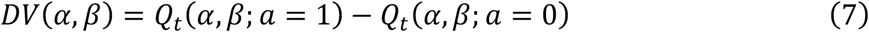

The optimal policy would be to sample when the DV is positive; however, we will also introduce decision noise temperature.

#### Decision noise

The model allows for decision noise through a logistic function:

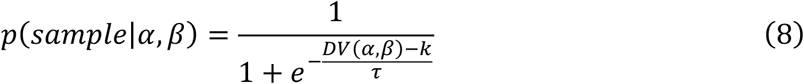

where *DV* is the decision variable in the model, *k* is a criterion parameter, and *τ* is the decision noise (higher *τ* means more noise). The noise is not part of the forward reasoning calculations. Instead, it is part of the decision to sample or stop once the optimal solution is derived.

#### Investment decisions with decision noise

For the models that use a Bayesian belief distribution (the CAS model, the Sample cost model and the Uncertianty model described below), the equation for the utility of investing is the same. Once the decision to stop sampling has been made, the agent choses to invest using the utility of investing with decision noise:

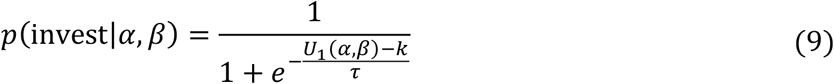

Note that we allow for a different temperature in the probability of sampling in all models.

## Acknowledgments

This study was funded by a European Research Council grant for A.G.S. (#313454). The funders had no role in study design, data collection and analysis, decision to publish or preparation of the manuscript. We thank Jeroen van Baar for helpful comments.

## References

1 Berg, J., Dickhaut, J. & McCabe, K. Trust, reciprocity, and social history. Games and economic behavior 10, 122–142 (1995).

2 Axelrod, R. & Dion, D. The further evolution of cooperation. Science 242, 1385–1390 (1988).

3 Vives, M.-L. & FeldmanHall, O. Tolerance to ambiguous uncertainty predicts prosocial behavior. Nature communications 9, 2156 (2018).

4 Jordan, J. J., Hoffman, M., Nowak, M. A. & Rand, D. G. Uncalculating cooperation is used to signal trustworthiness. Proceedings of the National Academy of Sciences 113, 8658–8663 (2016).

5 Capraro, V. & Kuilder, J. To know or not to know? Looking at payoffs signals selfish behavior, but it does not actually mean so. Journal of Behavioral and Experimental Economics 65, 79–84 (2016).

6 Fiedler, K. & Juslin, P. Information sampling and adaptive cognition. (Cambridge University Press, 2006).

7 Kappes, A. et al. Uncertainty about the impact of social decisions increases prosocial behaviour. Nature Human Behaviour 2, 573 (2018).

8 Bellman, R. On the theory of dynamic programming. Proceedings of the National Academy of Sciences 38, 716–719 (1952).

9 Chang, L. J., Doll, B. B., van’t Wout, M., Frank, M. J. & Sanfey, A. G. Seeing is believing: Trustworthiness as a dynamic belief. Cognitive psychology 61, 87–105 (2010).

10 Aimone, J. A., Houser, D. & Weber, B. in Proc. R. Soc. B. 20132127 (The Royal Society).

11 Simon, H. A. Theories of bounded rationality. Decision and organization 1, 161–176 (1972).

12 Gigerenzer, G. & Goldstein, D. G. Reasoning the fast and frugal way: models of bounded rationality. Psychological review 103, 650 (1996).

13 Ratcliff, R. A theory of memory retrieval. Psychological review 85, 59 (1978).

14 Drugowitsch, J., Moreno-Bote, R., Churchland, A. K., Shadlen, M. N. & Pouget, A. The cost of accumulating evidence in perceptual decision making. Journal of Neuroscience 32, 3612–3628 (2012).

15 Lewandowsky, S. & Farrell, S. Computational modeling in cognition: Principles and practice. (SAGE publications, 2010).

16 Hoffman, M., Yoeli, E. & Nowak, M. A. Cooperate without looking: Why we care what people think and not just what they do. Proceedings of the National Academy of Sciences 112, 1727–1732 (2015).

17 Hilbe, C., Hoffman, M. & Nowak, M. A. Cooperate without looking in a non-repeated game. Games 6, 458–472 (2015).

18 FeldmanHall, O. & Shenhav, A. Resolving uncertainty in a social world. Nature human behaviour, 1 (2019).

19 Camerer, C. F. Behavioral game theory: Experiments in strategic interaction. (Princeton University Press, 2011).

20 Sanfey, A. G. Social decision-making: insights from game theory and neuroscience. Science 318, 598–602 (2007).

21 Pérez-Escudero, A., Friedman, J. & Gore, J. Preferential interactions promote blind cooperation and informed defection. Proceedings of the National Academy of Sciences 113, 13995–14000 (2016).

22 Denrell, J. & March, J. G. Adaptation as information restriction: The hot stove effect. Organization Science 12, 523–538 (2001).

23 Baumeister, R. F., Bratslavsky, E., Finkenauer, C. & Vohs, K. D. Bad is stronger than good. Review of general psychology 5, 323 (2001).

24 Skowronski, J. J. & Carlston, D. E. Negativity and extremity biases in impression formation: A review of explanations. Psychological bulletin 105, 131 (1989).

25 Fiske, S. T. Attention and weight in person perception: The impact of negative and extreme behavior. Journal of personality and Social Psychology 38, 889 (1980).

26 Mende-Siedlecki, P., Baron, S. G. & Todorov, A. Diagnostic value underlies asymmetric updating of impressions in the morality and ability domains. Journal of Neuroscience 33, 19406–19415 (2013).

27 Wilson, R. K. & Eckel, C. C. Judging a book by its cover: Beauty and expectations in the trust game. Political Research Quarterly 59, 189–202 (2006).

28 Delgado, M. R., Frank, R. H. & Phelps, E. A. Perceptions of moral character modulate the neural systems of reward during the trust game. Nature neuroscience 8, 1611 (2005).

29 Fareri, D. S., Chang, L. J. & Delgado, M. R. Effects of direct social experience on trust decisions and neural reward circuitry. Frontiers in neuroscience 6, 148 (2012).

30 Clark, L., Robbins, T. W., Ersche, K. D. & Sahakian, B. J. Reflection impulsivity in current and former substance users. Biological psychiatry 60, 515–522 (2006).

31 Bradley, B. P., Mogg, K., Millar, N. & White, J. Selective processing of negative information: Effects of clinical anxiety, concurrent depression, and awareness. Journal of abnormal psychology 104, 532 (1995).

32 Hauser, T. U., Moutoussis, M., Dayan, P. & Dolan, R. J. Increased decision thresholds trigger extended information gathering across the compulsivity spectrum. Translational psychiatry 7, 1296 (2017).

33 Hauser, T. U. et al. Increased decision thresholds enhance information gathering performance in juvenile Obsessive-Compulsive Disorder (OCD). PLoS computational biology 13, e1005440 (2017).

34 King-Casas, B. & Chiu, P. H. Understanding interpersonal function in psychiatric illness through multiplayer economic games. Biological psychiatry 72, 119–125 (2012).

35 Lazarus, S. A., Cheavens, J. S., Festa, F. & Rosenthal, M. Z. Interpersonal functioning in borderline personality disorder: a systematic review of behavioral and laboratory-based assessments. Clinical Psychology Review 34, 193–205 (2014).

36 Hula, A., Vilares, I., Lohrenz, T., Dayan, P. & Montague, P. R. A model of risk and mental state shifts during social interaction. PLoS computational biology 14, e1005935 (2018).

37 Bohnet, I. & Zeckhauser, R. Trust, risk and betrayal. Journal of Economic Behavior & Organization 55, 467–484 (2004).

38 Bellman, R. A Markovian decision process. Journal of Mathematics and Mechanics, 679–684 (1957).

39 Mulder, M. J., Wagenmakers, E.-J., Ratcliff, R., Boekel, W. & Forstmann, B. U. Bias in the brain: a diffusion model analysis of prior probability and potential payoff. Journal of Neuroscience 32, 2335–2343 (2012).

40 Tajima, S., Drugowitsch, J. & Pouget, A. Optimal policy for value-based decisionmaking. Nature communications 7, 12400 (2016).

