## Supplementary Materials for "The Cost of Appearing Suspicious? Information Gathering in Trust Decisions"

**ONLINE SUPPLEMENTARY INFORMATION**

*IST trial procedure*

At the start of each trial, participants started with an investment endowment of 6 euros (E6) and a sampling budget of 25 x 5 eurocent. A trial started with the presentation of a starting screen that contained:

- A blurry photo of a person (the trustee). In the social context in which the trustee would not be informed of the sampling (overt sampling), a black bar partially covered the trustee's photo.
- A visualization of the condition: in the no cost condition, the word "free" was displayed, and in the cost condition, an image of tokens and a budget counter at 25.
- A 5-by-5 grid of empty boxes.
- A visualization and numerical representation of the initial counts of positive samples (green; 0), negative samples (red; 0), and unopened boxes (grey; 25).

The participant generally had three options: to click on an empty box, to choose "Invest E6", or to choose "Do not invest". If the number of unopened boxes was 0, then only the latter two options were available. There was no time limit for the decision. If the participant clicked on an empty square, its color was revealed. The color of the opened box was determined by a draw from a Bernoulli distribution with parameter equal to the reciprocation probability $r$ on that trial, $r$ (0, 0.2, 0.4, 0.6, 0.8, or 1). Simultaneously with the revealing of the box's color, the three counters were correspondingly updated, and in the monetary cost condition, 5 eurocents were subtracted from the sampling budget counter. The participant then made their next choice, still within the same trial. Once the participant clicked "Invest E6" or "Do not invest", a confirmation of their choice was shown for 1.5 seconds on an otherwise blank screen, and the next trial began. Outcomes of the investment decisions were not shown during the task.

*Descriptive statistics and mixed model*

Participants sampled on average of 11.5±0.5 times per trial (from a maximum of 25).

Descriptive statistics obtained using a mixed-effects model in R, package lme4 (www.r-project.org[1,2] R Core Team, 2014). The model included the interaction between monetary and social context conditions, the absolute difference between positive and negative sample outcomes, and a random intercept per participant.

**Heuristic computational models**

*Sample cost model*

This model is similar to the CAS model without weight parameter $\omega$. The utility of investing thus becomes:

$$U_1(\alpha, \beta) = \mathrm{E}[\text{outcome}|\alpha, \beta] - \lambda \mathrm{Var}[\text{outcome}|\alpha, \beta] = \frac{m\alpha}{\alpha + \beta} - \frac{\lambda m^2 \alpha\beta}{(\alpha + \beta)^2} \tag{10}$$

Instead of a decrease in the reciprocation probability *r*, there is a separately estimated cost parameter *c* for each condition (equation 6). The resulting policy is ran through a softmax to account for decision noise (equation 9).

*Uncertainty model*

The Uncertainty model is based on a criterion on the uncertainty about *r*. Similar to the CAS model, the prior over *r* is a beta distribution with parameters $\alpha_0$ and $\beta_0$ and the posterior over *r* is a beta distribution with parameters $\alpha = n_+ + \alpha_0$ and $\beta = n_- + \beta_0$ . The decision variable in the Uncertainty model is the standard deviation of this posterior distribution:

$$DV_{Uncertainty}(\alpha, \beta) = \sqrt{\frac{\alpha\beta}{(\alpha, \beta)^2(\alpha + \beta + 1)}} \tag{11}$$

Decision noise is again modelled using equation (9).

*Discrete Drift Diffusion model*

In the discrete Drift Diffusion model (*dDDM*), the agent keeps track of the absolute difference between positive and negative information and stops sampling when this difference reaches a bound. However, to be consistent with our other models and with most of the value-based decision literature, we will use soft rather than hard bounds.

The decision variable of the "soft"  is the absolute difference between these two variables:

$$DV_{\mathrm{dDDM}}(n_+, n_-) = |n_+ - n_-| \tag{12}$$

The probability that the agent stops sampling is a logistic function of the difference between this decision variable and a bound $b$:

$$p(\mathrm{sample}|n_+, n_-) = \frac{1}{1 + e^{-\frac{DV(n_+, n_-) - b}{\tau}}} \tag{13}$$

One can think of this model as a dDDM with decision noise. The parameter $\tau$ is the temperature of the decision noise. If $\tau$ approaches 0, the mapping from $DV_{\mathrm{dDDM}}$ to the decision becomes deterministic: stop sampling when $|n+-n-| > b$. The presence of decision noise means that the agent sometimes continues sampling even when $|n_+-n_-| > b$, and sometimes already stops sampling when this inequality is not met.

We consider three versions of the dDDM, which differ in the assumptions about the bound $b$. In the $\mathrm{dDDM_s}$ model (where "s" stands for "symmetric"), $b$ is a fixed constant. In the $\mathrm{dDDM_a}$ model (where "a" stands for "asymmetric"), $b$ takes three possible values, depending on the sign of $n_+-n_-$:

$$b = \begin{cases} b_+ & \text{if } n_+ > n_- \\ \frac{b_+ + b_-}{2} & \text{if } n_+ = n_- \\ b_- & \text{if } n_+ < n_- \end{cases} \tag{14}$$

In the absence of decision noise, this would be a dDDM with asymmetric bounds or with an initial offset[39]. One consequence of this model is that when the difference between positive and negative evidence is close to zero, this model will take a very long time to decide between investing or not investing. To prevent that the agent samples for too long, it may be optimal that the bounds of investing and not investing "collapse" towards zero over time[40]. We implemented the collapsing bounds, in the dDDMc model (where “c” stands for “collapsing”), by exponentially decaying the bounds towards zero with speed $s$ over time, where time is defined as the number of samples $n$:

$$b_c = b * e^{-s*n} \tag{15}$$

where $s$ is a free parameter. If $s$ is zero, this term is equivalent to a non-collapsing bound.

*Investment decisions*

Once the decision to stop sampling has been made, the agent choses to invest using the utility of investing with decision noise:

$$p(\text{invest}|\alpha,\beta) = \frac{1}{1 + e^{-\frac{n_+ - n_- - k_{\text{invest}}}{\tau_{\text{invest}}}}} \tag{16}$$

Note that we allow for a different temperature in the probability of sampling in all models.

*Fit improvement after additional free parameters in the heuristic models*

The basic Sample cost model (methods) has six free parameters; decision noise temperature, decision bias, and one parameter for the cost of a sample in monetary units in each of the four experimental conditions. In our paradigm, an individual’s initial belief about trustworthiness is a prior probability distribution over the probability of a reciprocation ($r$), before any information about the trustee is known. We modelled the prior as a beta distribution with two free parameters ($\alpha_0$ and $\beta_0$). We operationalized risk attitude by subtracting the variance of the outcome, multiplied by a free parameter $\lambda$, from the expected amount earned from investing to obtain the utility of the decision to invest (Methods). The introduction of the subjective prior yielded a large improvement over the basic model (Table S1, within model comparisons). The risk attitude parameter yielded further improvement (Table S1, within model comparisons). The median risk attitude parameter was estimated at 0.142 (bootstrapped 95% CI [0.092, 0.177]), which shows that people were overall risk-averse. The subjective prior beliefs and risk aversion account for the finding that people gather less information when $r = 0.2$ compared with $r = 0.8$ (Figure 1b): when initial sample outcomes are in favour of not investing, people stop gathering more evidence sooner than when the outcomes are in favour of investing. In the model, such an asymmetry arises from the fact that investing is risky and not investing is not risky: risk aversion leads to less interest in gathering information about a trustee who already appears untrustworthy.

In the *Uncertainty model*, the agent computes the posterior in the same way as the CAS and Sample cost model, but the subject samples until uncertainty – measured as the posterior standard deviation (Methods) – drops below a fixed criterion. The basic Uncertainty model has five free parameters: one decision noise parameter and one criterion parameter for each of the four experimental conditions. For the same reasons as described earlier, we tested the improvement in model fit when adding free priors. We found that allowing for a subjective prior improved the model fit. The median of individual prior means was $r = 0.59$ (bootstrapped 95% CI [0.56, 0.65]).

The basic version of the dDDM has five free parameters; decision noise temperature, and one bound for each of the four experimental conditions. Next, we sequentially examined two versions of the dDDM. First, we allowed for potential valence-dependent biases in information sampling[3-6] by examining the model fit improvement of the dDDM with asymmetric bounds for positive and negative samples (Methods). However, using separate bounds did not convincingly improve the model fit when using a more stringent punishment for the number of free parameters. Second, we tested the model when the bounds collapse to zero over time. This is a common approach in DDM's to avoid excessive sampling when there is high stochasticity in outcomes, and may be optimal[7,8]. The speed of this collapse is determined by a free parameter $s$ (Methods). Consistent with recently reviews on perceptual decision-making[9,10], we find that collapsing bounds did not improve the model fit when we used a more stringent punishment for the number of free parameters.

Table S1. Model comparisons study 1

| | AIC | | | BIC | | |
|---|---|---|---|---|---|---|
| | | 95% CI | | | 95% CI | |
| | median | lower | upper | median | lower | upper |
| **Within-model comparisons** | | | | | | |
| CAS basic vs. priors | 1779 | 1322 | 2261 | 1366 | 911 | 1848 |
| CAS priors vs. risk attitude | 969 | 420 | 1620 | 762 | 216 | 1415 |
| Sample cost basic vs. priors | 2006 | 1506 | 2550 | 1584 | 1085 | 2128 |
| Sample cost priors vs. risk attitude | 1062 | 615 | 1569 | 851 | 404 | 1360 |
| Uncertainty basic vs. priors | 1923 | 1211 | 2719 | 1501 | 786 | 2300 |
| dDDM basic vs. two bounds | 1005 | 463 | 1651 | 161 | -380 | 805 |
| dDDM two bounds vs. collapsing bound | 459 | 180 | 814 | 248 | -33 | 599 |
| **Between-models comparisons** | | | | | | |
| Sample cost vs. CAS | 825 | 70 | 1610 | 825 | 109 | 1725 |
| Uncertainty vs. CAS | -1268 | -2326 | 219 | -1681 | -2949 | -444 |
| Uncertainty vs. Sample cost | -2082 | -3597 | -635 | -2504 | -4023 | -1065 |
| Sample cost vs. dDDM | -3974 | -5239 | -2767 | -3130 | -4397 | -1920 |
| Uncertainty vs. dDDM | -6056 | -7055 | -5117 | -5634 | -6639 | -4689 |
| dDDM vs. CAS | 4786 | 3849 | 6175 | 3960 | 2787 | 5224 |

95% CI = Bootstrapped 95% confidence interval of the summed difference between model fits. Smaller AIC and BIC values indicate better fit. Thus, positive values indicate a better fit for the second model. The models were fitted to the data at the individual level using a log likelihood optimization algorithm as implemented in the fmincon routine in MATLAB (©Mathworks). The optimization was iterated 100 times with varying initiations to avoid local minima. Because of the summation of the difference, large positive or negative numbers therefore reflect that one model wins consistently, i.e. for most subjects.

*Condition effects on criterion parameter estimates for the heuristic models*

The condition-specific parameter estimates for the CAS model are mentioned in the main text. The condition-specific parameter estimates of the heuristic models are provided below.

Table S2. Between-condition estimates for the heuristic models study 1

| | Covert | | | Overt | | |
|---|---|---|---|---|---|---|
| | | 95% CI | | | 95% CI | |
| | median | lower | upper | median | lower | upper |
| Sample cost model *c* no cost | 0.000 | 0.000 | 0.017 | 0.037 | 0.008 | 0.165 |
| Sample cost model *c* cost | 0.716 | 0.171 | 1.130 | 0.758 | 0.214 | 1.310 |
| Uncertainty model *k* no cost | 0.028 | 0.022 | 0.053 | 0.075 | 0.041 | 0.092 |
| Uncertainty model *k* cost | 0.119 | 0.105 | 0.131 | 0.122 | 0.112 | 0.129 |
| Drift Diffusion model *b* no cost | 18.205 | 15.109 | 25.846 | 12.432 | 7.691 | 21.985 |
| Drift Diffusion model *b* cost | 8.271 | 7.294 | 9.815 | 8.647 | 7.326 | 10.288 |

Bootstrapped 95% confidence interval of the median. Estimated cost parameters of the Sample cost model are normalized on the objective cost of 5 euro cents. Cost of a sample (*c*), stopping criterion (*k*), decision bound (*b*).

Table S3. Free parameters per model

| **Model** | **Global parameters** | **Parameters per cost condition** |
|---|---|---|
| CAS | $\tau$, $k_0$ | $c$ (cost and no cost), $\omega$ (overt cost, overt no cost) |
| CAS priors | $\tau$, $k_0$, $\alpha_0$, $\beta_0$ | $c$ (cost and no cost), $\omega$ (overt cost, overt no cost) |
| CAS priors and risk attitude | $\tau$, $k_0$, $\alpha_0$, $\beta_0$, $\lambda$ | $c$ (cost and no cost), $\omega$ (overt cost, overt no cost) |
| $dDDM_a$ | $\tau$ | $b$ |
| $dDDM_s$ | $\tau$ | $b_+$, $b_-$ |
| $dDDM_c$ | $\tau$, $s$ | $b$ |
| Sample cost | $\tau$, $k_0$ | $c$ |
| Sample cost priors | $\tau$, $k_0$, $\alpha_0$, $\beta_0$ | $c$ |
| Sample cost priors, risk attitude | $\tau$, $k_0$, $\alpha_0$, $\beta_0$, $\lambda$ | $c$ |
| Uncertainty | $\tau$ | $k$ |
| Uncertainty priors | $\tau$, $\alpha_0$, $\beta_0$ | $k$ |

$\tau$ = Decision noise temperature, = $k_0$=softmax intercept, $\lambda$ = risk attitude*,* $\alpha_0$ = alpha prior*,* $\beta_0$ = beta prior, $c$ = sampling cost, $k$ = stopping criterion, $b$ = decision bound. Through parameter recovery, we verified that the number of trials was large enough to accurately estimate parameters.

To test if the model predictions could be distinguished from each other and test for biases against any particular model, we performed model recovery. We used the parameter estimates to generate data from each model, and fitted those data to each model. This shows that all three models were recoverable, as data generated by a model was also best fitted by that model (Table S3).

Table S4. Model recovery results

| | AIC | | | BIC | | |
|---|---|---|---|---|---|---|
| | | 95% CI | | | 95% CI | |
| | median | lower | upper | median | lower | upper |
| **Data generated by the CAS model** | | | | | | |
| CAS vs. Sample cost | -1947 | -2514 | -1396 | -1947 | -2585 | -1367 |
| CAS vs. Uncertainty | -2909 | -3473 | -2400 | -2679 | -3233 | -2199 |
| CAS vs. dDDM | -2145 | -3559 | -639 | -1685 | -3185 | -319 |
| **Data generated by the Sample cost model** | | | | | | |
| Sample cost vs. CAS | -388 | -702 | -54 | -388 | -738 | -109 |
| Sample cost vs. Uncertainty | -2758 | -3176 | -2175 | -2516 | -3060 | -1980 |
| Sample cost vs. dDDM | -3833 | -4673 | -2972 | -3349 | -4273 | -2466 |
| **Data generated by the Uncertainty model** | | | | | | |
| Uncertainty vs. CAS | -2681 | -3634 | -1709 | -2912 | -3888 | -2018 |
| Uncertainty vs. Sample cost | -2581 | -3710 | -1657 | -2811 | -3740 | -1978 |
| Uncertainty vs. dDDM | -3210 | -4187 | -2474 | -2402 | -3356 | -1674 |
| **Data generated by the dDDM** | | | | | | |
| dDDM vs. CAS | -1168 | -1658 | -703 | -2201 | -2754 | -1667 |
| dDDM vs. Sample cost | -1654 | -2140 | -1165 | -2114 | -2641 | -1612 |
| dDDM vs. Uncertainty | -1723 | -2383 | -1164 | -1953 | -2523 | -1424 |

95% CI = Bootstrapped 95% confidence interval of the summed difference between model fits. Negative values indicate a better fit of the model that generated the data. The data were generated using the participants' parameter estimates. This shows that all models were recoverable.

We further examined models could also predict investment decisions (after sampling has concluded) by using the parameter estimates. We therefore extracted the expected utility of investing at the time of stopping for the Sample cost model and for the Uncertainty model and fitted those expected values to the decisions to invest or not invest using a logistic regression. For the dDDM, we took the difference between the positive and negative sample outcomes as predictor variable in the logistic regression. These regressions allowed for bias and decision noise temperature (equation 9). The fits of the Sample cost and Uncertainty models were not significantly different (95% CI of the summed difference in log likelihoods [-54.66, 46.44]) but the dDDM performed less well. Both models resulted in a significant predictor of the probability of investing (Uncertainty model $p < 0.001$, Nagelkerke pseudo-$R^2 = 0.882$; Sample cost model $p < 0.001$, pseudo-$R^2 = 0.832$; dDDM, $p < 0.001$, pseudo-$R^2 = 0.733$, Figure S1).

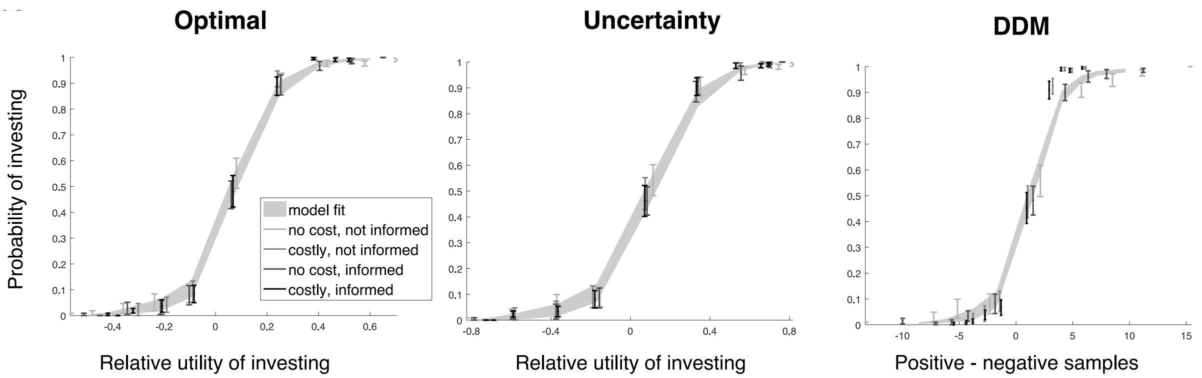


Figure S1. Model fits to the invest decisions for each model using their decision variables. For the Sample cost and Uncertainty models the decisions to invest are plotted as a function of expected utility of investing derived from the models. For the dDDM, the decision variable is the difference between positive and negative samples. Error bars represent the data (mean) ± SEM across all subjects. Shaded area is the model fit ± SEM.

***Study 2***

Each subject completed a total of 200 trials (40 trials per *r*), resulting in a maximum of 1000 sampling decisions per probability. There were 42 subjects in the positively biased condition, the data from 3 participants were excluded because they reported confusion about the task, resulting in 39 subjects (14 males, age: *m* = 23.15, *sd* = 2.75, range = 18-34 years). There were 39 subjects in the negatively biased condition, data from 2 participants were excluded because they did not understand the instructions, and data from 1 participant was excluded due to a technical malfunction, resulting in 36 subjects (11 males, age: *m* =21.16, *sd* = 3.13 range = 18-34 years). A one-way ANOVA showed no significant age difference between any of the groups in Study 1 and Study 2 ($p$ = .091).

*Descriptive statistics Study 2*

Mixed-effects models using the R package lme4 replicated all results of the sampling decision analyses in study 1. The mixed model included monetary cost, social context, bias, and outcome uncertainty, their interactions and a random intercept for each participant. We used a separate logistic regression to test whether the decision to invest was predicted by $r$ and the cost conditions. This also replicated our findings, as the regression returned 2.989±0.048, $p < 0.001$ for $r$, indicating that the probability of investing increased with a higher $r$. This confirms that the acquired information was actually used in the final investment decision. As expected, the two cost manipulations (monetary and social) and their interaction were not significant predictors of the decision to invest (monetary cost: -0.031±0.027, $p$ = 0.248; social context: -0.045±0.027, $p$ = 0.091; interaction between monetary and social context: 0.047±0.027, $p > 0.081$).

Table S5. Model comparisons Study 2

| | **AIC** | | | **BIC** | | |
|---|---|---|---|---|---|---|
| | | 95% CI | | | 95% CI | |
| | median | lower | upper | median | lower | upper |
| **Within-models comparisons** | | | | | | |
| CAS basic - priors | 3273 | 2678 | 3899 | 2409 | 1812 | 3033 |
| CAS priors – risk attitude | 2385 | 1515 | 3407 | 1953 | 1082 | 2982 |
| Sample cost basic – priors | 1755 | 1184 | 2367 | 891 | 322 | 1504 |
| Sample cost priors - risk attitude | 5442 | 4142 | 6862 | 5010 | 3714 | 6426 |
| Uncertainty basic - priors | 6145 | 3925 | 8855 | 5281 | 3072 | 7983 |
| dDDM basic – two bounds | 6561 | 3388 | 9684 | 4833 | 1657 | 7977 |
| dDDM two bounds – collapsing bound | 2709 | -3522 | 7486 | 2277 | -3957 | 7026 |
| **Between-models comparisons** | | | | | | |
| Sample cost – CAS | 794 | -336 | 1783 | 794 | -336 | 1783 |
| Uncertainty - CAS | 1032 | -756 | 2271 | 168 | -1440 | 1908 |
| Uncertainty - Sample cost | -238 | -2365 | 1731 | 626 | -1498 | 2600 |
| Sample cost – dDDM | -6761 | -8373 | -5319 | -2873 | -4505 | -1417 |
| Uncertainty - dDDM | -3500 | -4908 | -2006 | -3500 | -4893 | -2024 |
| dDDM - CAS | 5395 | 4110 | 6709 | 3668 | 2383 | 4988 |

95% CI = Bootstrapped 95% confidence interval of the summed difference between model fits. Because of the summation of the difference, large positive or negative numbers therefore reflect that one model wins consistently, i.e. for most subjects. Lower AIC and BIC values indicate a better model fit. Thus, positive values indicate a better fit for the second model, while negative values indicate a better fit for the first model. The models were fitted using an identical procedure to that described in study 1. The distribution of generative reciprocation probabilities was biased in study 2. For the basic CAS, Sample cost, and Uncertainty models, we therefore fixed the priors based on the true distribution (alpha = 0.84 and beta = 0.56) for the positively biased distribution and vice versa for negatively biased distribution.

*Parameter estimates Study 2*

The Cost of Appearing Suspicious $\omega$ estimate was higher when sampling was overt, replicating the social context effect ($Z = -3.77$, $p < 0.001$). The sampling cost $c$ was higher when sampling was monetarily costly compared to when it was cheap ($Z$ = -5.169, $p$ < 0.001). Risk attitude estimates again suggested that people were overall risk averse (median = 0.037 95% CI [-0.060, 0.128]).

Table S6. Median estimated condition parameters for heuristic models Study 2

| **Positive Bias** | Not informed | | | Informed | | |
|---|---|---|---|---|---|---|
| | | 95% CI | | | 95% CI | |
| | median | lower | upper | median | lower | upper |
| Sample cost model $c$ no cost | 0.003 | 0.000 | 0.024 | 0.025 | 0.012 | 0.205 |
| Sample cost model $c$ cost | 1.059 | 0.233 | 2.305 | 1.125 | 0.527 | 2.191 |
| Uncertainty model $k$ no cost | 0.052 | 0.030 | 0.061 | 0.062 | 0.048 | 0.073 |
| Uncertainty model $k$ cost | 0.100 | 0.084 | 0.112 | 0.101 | 0.089 | 0.113 |
| dDDM $b_+$ no cost | 21.214 | 17.632 | 28.182 | 18.230 | 10.705 | 21.392 |
| dDDM $b_+$ cost | 9.355 | 7.698 | 11.860 | 9.803 | 7.325 | 12.403 |
| dDDM $b_-$ no cost | 16.056 | 12.738 | 19.984 | 8.342 | 6.691 | 10.411 |
| dDDM $b_-$ cost | 12.648 | 9.103 | 16.720 | 7.471 | 6.355 | 9.928 |
| **Negative bias** | | | | | | |
| Sample cost model $c$ no cost | 0.000 | 0.000 | 0.018 | 0.255 | 0.013 | 0.550 |
| Sample cost model $c$ cost | 0.787 | 0.309 | 1.231 | 0.726 | 0.3845 | 1.344 |
| Uncertainty model $k$ no cost | 0.014 | 0.00 | 0.028 | 0.057 | 0.023 | 0.074 |
| Uncertainty model $k$ cost | 0.108 | 0.094 | 0.114 | 0.106 | 0.094 | 0.125 |
| dDDM $b_+$ no cost | 19.584 | 15.372 | 27.190 | 12.102 | 9.458 | 23.682 |
| dDDM $b_+$ cost | 9.599 | 7.211 | 11.859 | 9.526 | 6.945 | 11.868 |
| dDDM $b_-$ no cost | 20.961 | 15.339 | 24.669 | 9.599 | 7.641 | 11.091 |
| dDDM $b_-$ cost | 14.369 | 9.984 | 20.188 | 9.274 | 7.207 | 10.899 |

95% CI = Bootstrapped 95% confidence interval of the median. Estimated cost parameters of the Sample cost model are normalized on the objective cost of 5 euro cents. Cost of a sample ($c$), stopping criterion ($k$), decision bound ($b$).

The expected utility of investing derived from the CAS model fitted the invest decisions ($\beta$ = 3.720, $p$ <0.001, Nagelkerke pseudo-$R^2$ = 0.40). The Sample cost model showed a weaker fit ($p$ < .001, Nagelkerke pseudo-$R^2$ = 0.15), while the expected utility of investing derived from the

Uncertainty model showed a stronger relation with investing ($p < .001$, Nagelkerke pseudo-$R^2$ = 0.64), as did the dDDM (Nagelkerke pseudo-$R^2$ = 0.69, $p < .001$).

*Subjective reports Study 2*

The subjective reports in study 2 replicated the pattern in study 1: Over half of the participants believed that the reciprocation probability $r$ would decrease when information was overtly sampled (51% and 59% in positive and negative bias conditions, respectively). Fewer believed that the probability would become larger (27% and 15% in the positive and negative bias conditions, respectively). The remaining participants believed it would not affect $r$.

**References**


1 Bates, D., Mächler, M., Bolker, B. & Walker, S. Fitting linear mixed-effects models using lme4. *arXiv preprint arXiv:1406.5823* (2014).

2 Kuznetsova, A., Brockhoff, P. B. & Christensen, R. H. B. lmerTest package: tests in linear mixed effects models. *Journal of Statistical Software* **82** (2017).

3 Baumeister, R. F., Bratslavsky, E., Finkenauer, C. & Vohs, K. D. Bad is stronger than good. *Review of general psychology* **5**, 323 (2001).

4 Reeder, G. D. & Coovert, M. D. Revising an impression of morality. *Social Cognition* **4**, 1-17 (1986).

5 Siegel, J. Z., Mathys, C., Rutledge, R. B. & Crockett, M. J. Beliefs about bad people are volatile. *Nature Human Behaviour* **2**, 750 (2018).

6 Skowronski, J. J. & Carlston, D. E. Negativity and extremity biases in impression formation: A review of explanations. *Psychological bulletin* **105**, 131 (1989).

7 Drugowitsch, J., Moreno-Bote, R., Churchland, A. K., Shadlen, M. N. & Pouget, A. The cost of accumulating evidence in perceptual decision making. *Journal of Neuroscience* **32**, 3612-3628 (2012).

8 Tajima, S., Drugowitsch, J. & Pouget, A. Optimal policy for value-based decision-making. *Nature communications* **7**, 12400 (2016).

9 Hawkins, G. E., Forstmann, B. U., Wagenmakers, E.-J., Ratcliff, R. & Brown, S. D. Revisiting the evidence for collapsing boundaries and urgency signals in perceptual decision-making. *Journal of Neuroscience* **35**, 2476-2484 (2015).

10 Ratcliff, R., Smith, P. L., Brown, S. D. & McKoon, G. Diffusion decision model: current issues and history. *Trends in cognitive sciences* **20**, 260-281 (2016).
